# Rainfall as a significant contributing factor to butterfly seasonality along a climatic gradient in the neotropics

**DOI:** 10.1101/630947

**Authors:** María F. Checa, Elisa Levy, Jaqueline Rodriguez, Keith Willmott

## Abstract

We analyzed the dynamics of multi-species butterfly communities along a climatic gradient with varying precipitation regimes for three consecutive years, and determine how climatic variables associate with observed butterfly seasonality. To provide a baseline for future studies of how climate change might affect these butterfly populations, we additionally explored the role of butterfly seasonality as a potential contributing factor for their susceptibility to climate variation. As far as we know, this represents the first study that simultaneously sampled and described seasonality patterns of tropical butterfly communities across ecosystems with varying climatic seasonality. A 3-year survey was carried out at three sites (i.e., wet, transition and dry forests) across a climatic gradient in western Ecuador. Butterflies were sampled using traps baited with rotting banana and prawn every two months from Nov 2010 to Sep 2013. Traps were set up at two heights, in the understory and canopy. In total, 7046 individuals of 212 species were sampled over 180 sampling days.

Butterfly communities exhibited conspicuous intra and inter-annual variation in temporal dynamics with certain elements (e.g., maximum abundance recorded) of seasonality patterns likely synchronized in seasonal forests (i.e., transition and dry forest) across years but not in aseasonal forests (i.e., wet forest). In addition, the highest numbers of species and individuals occurred during the wet season across all study sites and years; indeed, rainfall was significantly positively associated with temporal abundance. Likewise, butterfly species displaying stronger seasonality were significantly associated with higher rainfall periods in seasonal forests. Variation in precipitation regimes might significantly affect more seasonal species.

## Introduction

The mechanisms driving higher biodiversity towards the equator have puzzled ecologists for decades. Recently, studies into insects have contributed to our understanding of these mechanisms by focusing on host-specificity in insect herbivores and spatial turnover (Dyer *et al.* 2007, Novotny *et al.* 2007). A further, little explored contributing factor might be the temporal partitioning of resources, thereby decreasing competition and promoting species coexistence (Grimbacher *et al.* 2009). For most insects a ‘temporal window’ exists when environmental conditions are most favorable for certain stages of their life cycle; synchronization of critical stages of the life cycle with this temporal window is thus expected to prevent fitness consequences (van Asch & Visser 2007). For example, numerous herbivores use specific plant resources during short periods of time, when the quality of these resources is optimal (Hellmann 2002), and as a consequence a peak in insect herbivore abundance is expected to coincide with leaf flush (Murakami *et al.* 2008, Srygley *et al.* 2010) or other resource availability (*e.g.*, rotten fruit, flowers) (Hamer *et al.* 2006, Checa *et al.* 2009). Competition for such resources could therefore drive temporal niche partitioning.

The population dynamics of insects in the form of seasonality patterns is one potential mechanism by which species might partition resources. For decades, it has been known that some tropical insects have marked seasonal changes in their relative abundance with climate as a key factor controlling their population dynamics (Wolda 1988, see review Kishimoto-Yamada & Itioka 2015). A pattern repeatedly found in the majority of studies of the temporal dynamics of tropical insects is an abundance peak during the wet season (Wolda 1978, Novotny & Basset 1998, Grimbacher *et al.* 2009 and citations therein), whereas other studies have found abundance to be higher during the dry season (see Valtonen *et al.* 2013). In addition, opposing patterns might occur even at local scales. For example, in one study, the spatiotemporal dynamics of insects differed in moist/shaded sites versus drier sites (Janzen 1973, Richards & Windsor 2007), with insect abundance reaching a peak in the wet season in the former, but peaking in the dry season in the latter. Butterfly data apparently support these results at regional scales. In Neotropical dry forests, butterfly abundance peaks during the months with highest precipitation and relative humidity in Mexico (Pozo *et al.* 2008, Torres *et al.* 2009), Venezuela (Shahabuddin *&* Terborgh 1999), and western Ecuador with relative humidity, but not temperature as a significant associated factor with this temporal abundance variation (Checa 2010, Checa *et al.* 2014). In contrast, butterfly communities of relatively aseasonal or rainforests in the Neotropics may show decreased species richness and abundance during the wet season but peaks through the transitional months (see DeVries & Walla 2001, Checa *et al.* 2009, Grøtan *et al.* 2012, Valtonen *et al.* 2013, Grøtan *et al.* 2014) or warmest part of the year (Ribeiro *et al.* 2010). Conversely, annual fluctuation of butterfly numbers are significantly and solely correlated with temperature within these ecosystems (Checa *et al.* 2009, Ribeiro *et al.* 2010, Grøtan *et al.* 2014). It is thus possible that temporal patterns of insect abundance and their associated climatic factors diverge across forests with varying seasonality. Further research is therefore needed to better understand the role of climate on insect communities from different biome types. The role of environmental factors influencing insect seasonality in the tropics is still poorly understood due to the challenges involved in maintaining long-term studies covering multiple years and generations (Azerefegne *et al.* 2001, Bonebrake *et al.* 2010, Valtonen *et al.* 2013, Kishimoto-Yamada & Itioka 2015).

Improved understanding of insect seasonality and underlying factors is valuable for identifying how insects will respond or adapt to climate change (Valtonen *et al.* 2013). Climate change will increase rainfall seasonality, which might increase the length or severity of the dry season, and drought incidence, as a consequence of El Niño–Southern Oscillation (ENSO) (Walsh & Ryan 2000). Hence, studies that analyze seasonality and species richness patterns in Western Ecuador, an area directly affected by ENSO could provide more insights about possible effects of seasonality changes of rainfall in the biota. This information might be in turn useful to determine the susceptibility to climate variability of butterfly species displaying different traits. Body size, population density and mobility contribute to extinction vulnerability of taxa (Brown 1971, Kattan *et al.* 1994, Shahabuddin & Ponte 2005, Graves & Gotelli 1983). Butterfly seasonality, however, might be an additional important and hitherto unstudied contributing factor as more seasonal species can have a more restricted ecological niche, which in turn might make them more vulnerable to climate and habitat change.

Western Ecuador offers an ideal opportunity to examine how the dynamics of multi-species butterfly communities from seasonal and aseasonal forests are related to climate, because it presents a marked gradient of life zones, with wet forests dominating in the north, which gradually change to moist forest and dry forest in the south (Dodson & Gentry 1991). Here, we analyze the dynamics of multi-species butterfly communities along a climatic gradient with varying precipitation regimes for three consecutive years, and determine how climatic variables associate with observed butterfly seasonality. To provide a baseline for future studies of how climate change might affect these butterfly populations, we additionally explored the role of butterfly seasonality as a potential contributing factor for their susceptibility to climate variation.

## Methods

### Census Techniques

Butterflies were sampled using baited-traps at three study sties: Canande River Reserve (CR, wet forest), Lalo Loor Dry Forest (LLDFR, transition forest) and Jorupe Reserve (JR, dry forest). Sampling was performed every two months (*i.e.*, 6 times a year) over 7 days each sampling period for three consecutive years from November 2010 to September 2013.

Van Someren-Rydon bait traps were used with two different types of baits: rotting shrimp fermented for 13-18 days, and banana fermented for 2 days. Traps were checked daily during the first 7 days of the sampling month from 9 am to 3 pm; traps were opened and baited on the first trapping day, and checked over the next 6 days with trapped butterflies being collected, except for the most abundant species, which were marked and released.

Two transects were established at each reserve with each transect containing eight sampling positions, with a minimum distance between sampling positions of 40 m. Two baited traps were set up at each sampling position in two different strata, understory (1.5 meters above the ground) and canopy (10-25 m depending on the ecosystem sampled). The use of banana and shrimp baits alternated between positions, thus neighboring positions had different types of bait. Canopy and understory traps in the same position were baited with the same type of bait. The order at which sampling positions were visited was modified every sampling day in order to correct any bias produced by both daily temporal (Wolda 1988, Sutcliffe *et al.* 1996) and spatial distribution of butterflies at each sampling point.

Data loggers were deployed at each reserve to register daily rainfall, temperature and relative humidity, and climatic data for the entire sampling period (*i.e.*, from Nov 2010 to Sept 2013) was thus obtained for subsequent analyses.

All collected material was examined and identified to species, and classified following the higher taxonomic classification of Wahlberg *et al.* (2009). Moreover, butterfly species were further classified as wet or dry species. Due to limited published data about geographic distribution of most species included in the present study, a qualitative approach was used for this classification. Butterfly species recorded at CRR (wet forest) but not at JR (dry forest) were considered as wet species, whereas species registered at JR but not at CRR were classified as dry species. All collected specimens were deposited in the Museum of Invertebrates of the Pontificia Universidad Católica del Ecuador (QCAZ), and the McGuire Center for Lepidoptera and Biodiversity, Florida Museum of Natural History at the University of Florida (MGCL).

### Statistical Analyses

Generalized linear models were used to test whether changes in both butterfly species richness and abundance were associated with local climatic variables (*i.e.*, rainfall, temperature and relative humidity) across study sites. Generalized linear models have applications in circumstances where the assumptions of the standard linear model do not hold, as in the case where the distribution of the data is not normal (Littell *et al.* 1996) or when data are autocorrelated in terms of space or time. Hence, generalized linear models permit analyses of response variables such as counts, binary, proportions and positive valued continuous distributions (Hilbe 1994, Hoffman 2004). Typically, count data do not follow a normal distribution, but a Poisson or a negative binomial. Key assumptions of GLN are homogeneity, normality and independence of residuals (Dobson 2002, Hoffman 2004).

Two different models were performed to analyze each dependent variable, butterfly species richness and abundance per study site. In order to perform these analyses, data from different strata and transects collected during a single month were pooled together. Each point in the analyses thus consisted of the number of butterflies collected in a given month, regardless of collection point (*i.e.*, strata, sampling position and transect), and the associated climatic variables registered for that specific month. Best fit models were selected using their values of the Akaike’s Information Criterion (AIC) and residual variance. Scatterplots of the residuals and fitted values were checked to examine whether the model met the assumption of homogeneity, in addition to normal probability plots of residuals to verify normal distribution of residuals (see Lindsey 1997, Dobson 2002). The negative binomial distribution fitted the data better than a Poisson or normal distribution, and the best models included total rainfall and mean temperature. Therefore, further interpretation is based on the outputs of these models.

Secondly, I investigated whether butterflies that were more seasonal, showing more restricted emergence peaks, were associated with any particular climatic variables, in the JR dry forest. As a measure of seasonality I used the coefficient of variation (CV) of abundance, calculated by dividing the standard deviation of log-transformed species abundance in each of the 18 consecutive months over the average log-transformed abundance of the 18 months. Abundance data were log-transformed to correct bias induced by species abundance. The CV constitutes a useful measure of seasonality or temporal heterogeneity of species distributions (see Hanya *et al.* 2011, Tello & Stevens 2010), and rather than reflecting seasonality patterns (*i.e.*, timing or length of species occurrence), it measures the evenness of the temporal distribution of species over sampling time. Higher values indicate broader variability of species occurrence whereas low values reveal a more even distribution. To measure the climatic niche when species were active as adults, the climatic variables of the months when species were recorded were selected and averaged. Climatic variables included total monthly rainfall, maximum and minimum values of relative humidity and temperature, and standard deviation of both temperature and relative humidity. The climatic niche measured corresponds to the realized climatic niche and not the fundamental niche (see Schweiger *et al.* 2014). Finally, I used General Linear Models to test whether climate (explanatory variables) explained seasonality (response variable) across butterfly species. The GLM excluded singletons and doubletons to improve fit. Best fit models were based on the Gaussian distribution, and included total monthly rainfall, and the standard deviation of both relative humidity and temperature.

## Results

In total, 7046 individuals of 212 species were sampled over 180 sampling days, representing the families Nymphalidae (Apaturinae, Biblidinae, Charaxinae, Cyrestinae, Danainae, Heliconiinae, Limenitidinae, Nymphalinae and Satyrinae), Hesperiidae, Riodinidae and Lycaenidae. Observed species richness was two-fold higher in the wet forest (129) compared to the dry forest (57), even though nearly three times greater butterfly abundance was recorded in the latter (3611). The transition forest showed intermediate numbers of species richness (91) and abundance (2039). Diversity and evenness of species abundance therefore decreased from wet forests towards dry forests in the south.

The climate of the study sites was distinctly seasonal (Fig. 1). Most of the rainfall occurred during the wet season from January to May, but was nearly absent in the transition and dry forest after August. The wet forest received considerably more precipitation throughout the study duration with annual cumulative values constantly exceeding 3,000 mm^3^, a two- and three-fold increase compared with the annual precipitation in the transition and dry forests, respectively. Furthermore, cumulative rain achieved highest values in 2012 ranging from 3886 mm^3^ in the wet forest to 1203 mm^3^ in the dry forest. Mean monthly temperature fluctuated by as much as 2-3°C from the warmest to the coldest months across years and reserves, although temperature varied more and reached higher maximum values in LLDFR (transition) and JR (dry forest) (Fig. 1).

**Figure 1.**
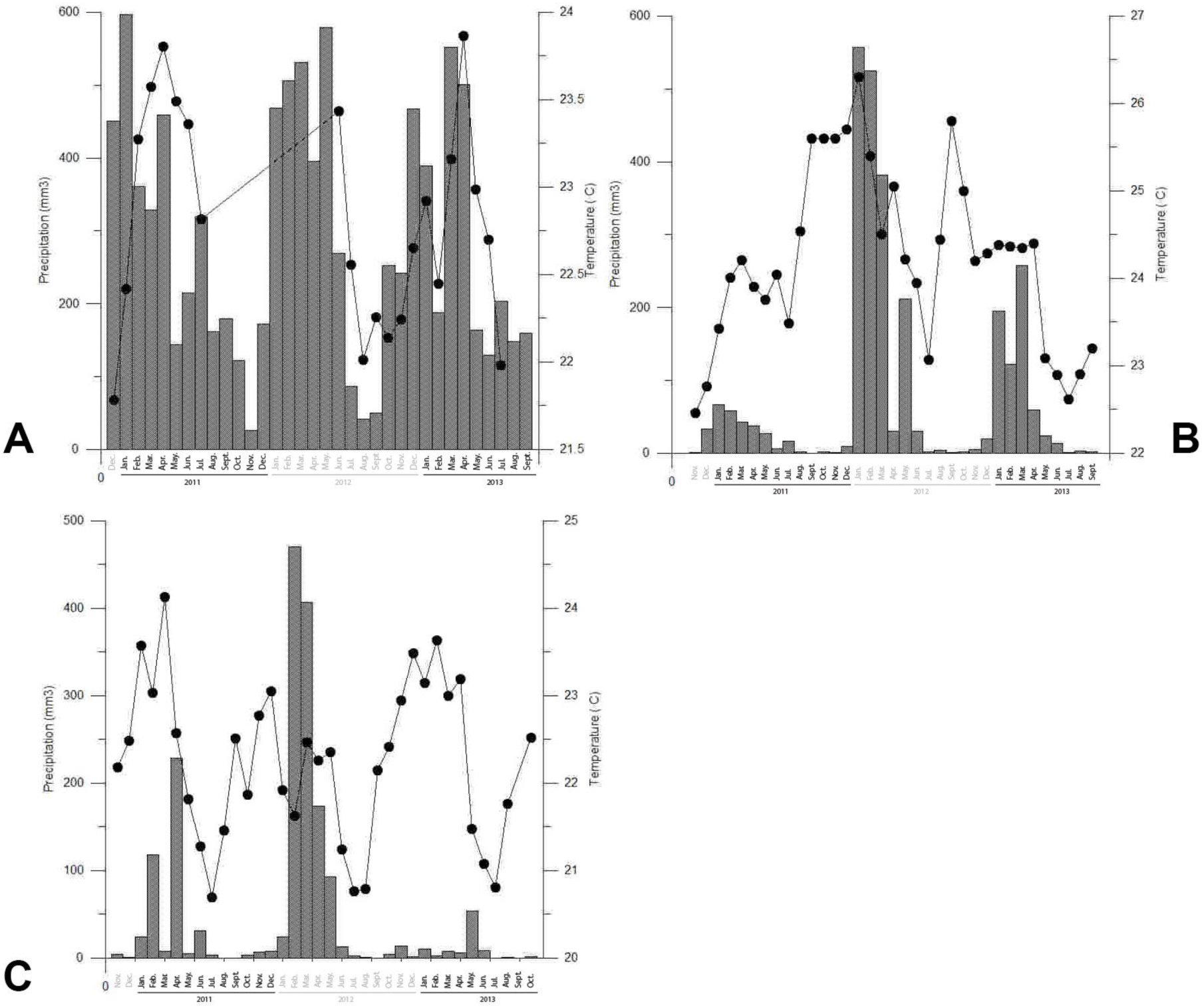
Monthly average temperature (line with dots) and monthly total rainfall (bars) recorded along a climatic gradient in Western Ecuador from November 2010 to September 2013. A) Wet forest located in the northern province of Esmeraldas (Canande River Reserve, CRR), B) Transition forest within Manabi province (Lalo Loor Dry Forest Reserve LLDFR) and C) Dry forest in the southern corner (Jorupe Reserve JR).

Butterfly communities showed conspicuous intra- and inter-annual variation in temporal dynamics with important similarities between the seasonality patterns of seasonal forests (*i.e.*, both transition and dry forests) compared to aseasonal forests (*i.e.*, wet forest) (Fig. 2). Furthermore, the greatest species richness and abundance were recorded during the wettest part of the year across all study sites. In both wet and transition forests, butterfly numbers reached a peak in March for both 2011 and 2013, and in May of 2012 (Fig. 2).

**Figure 2.**
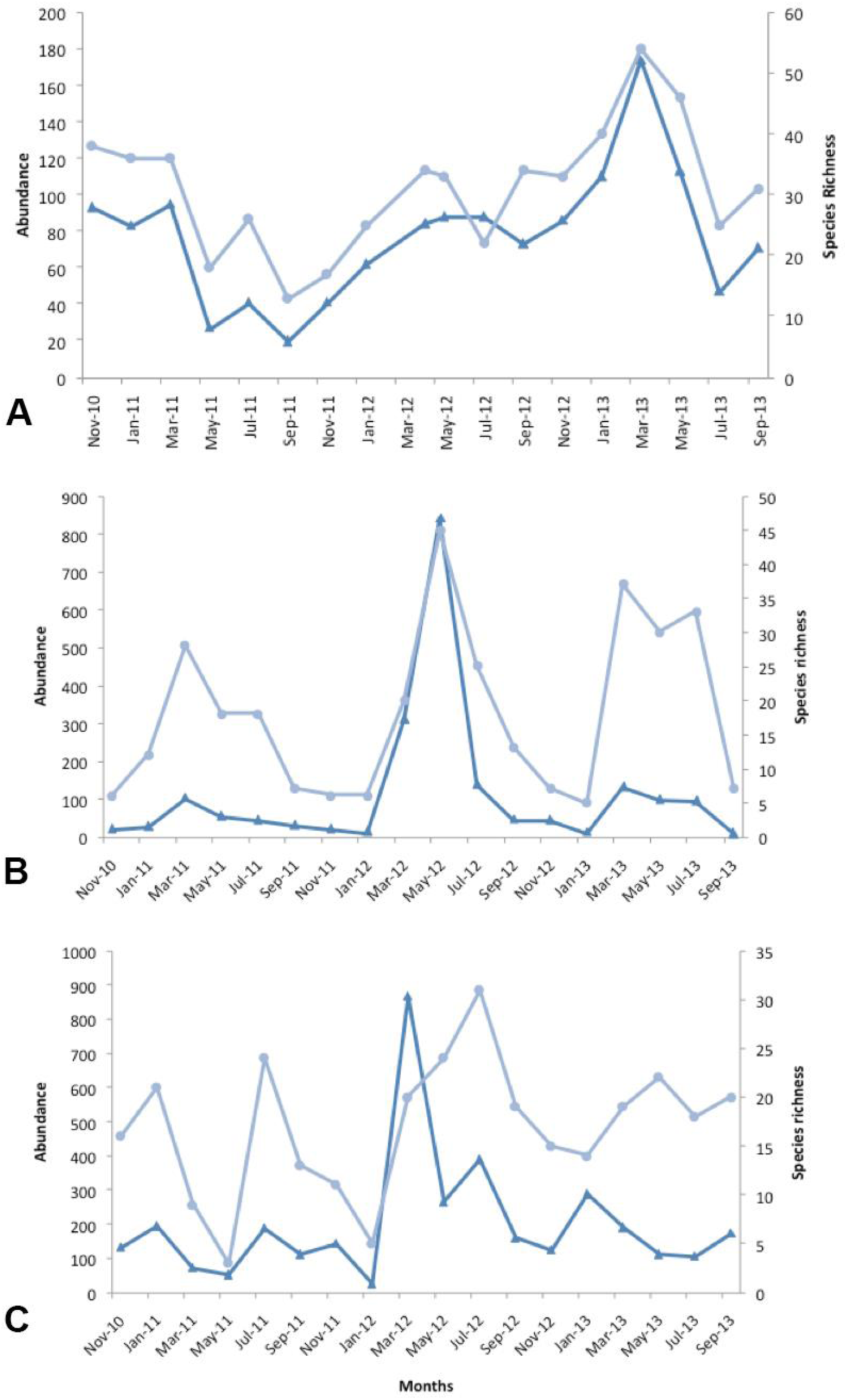
Monthly observed butterfly species richness (line with dots) and abundance (line with triangles) from Nov 2010 to Sep 2013 across study sites. A) wet forest, B) transition forest and C) dry forest.

In contrast, the highest numbers of species and individuals were recorded in January (2011 and 2013) and March (2012) in the dry forest. A second, weaker peak occurred in July and/or September across all study sites over all years. These temporal patterns remained similar for remaining species when extracting most abundant species (>100 individuals) from plots (Fig. 3).

**Figure 3.**
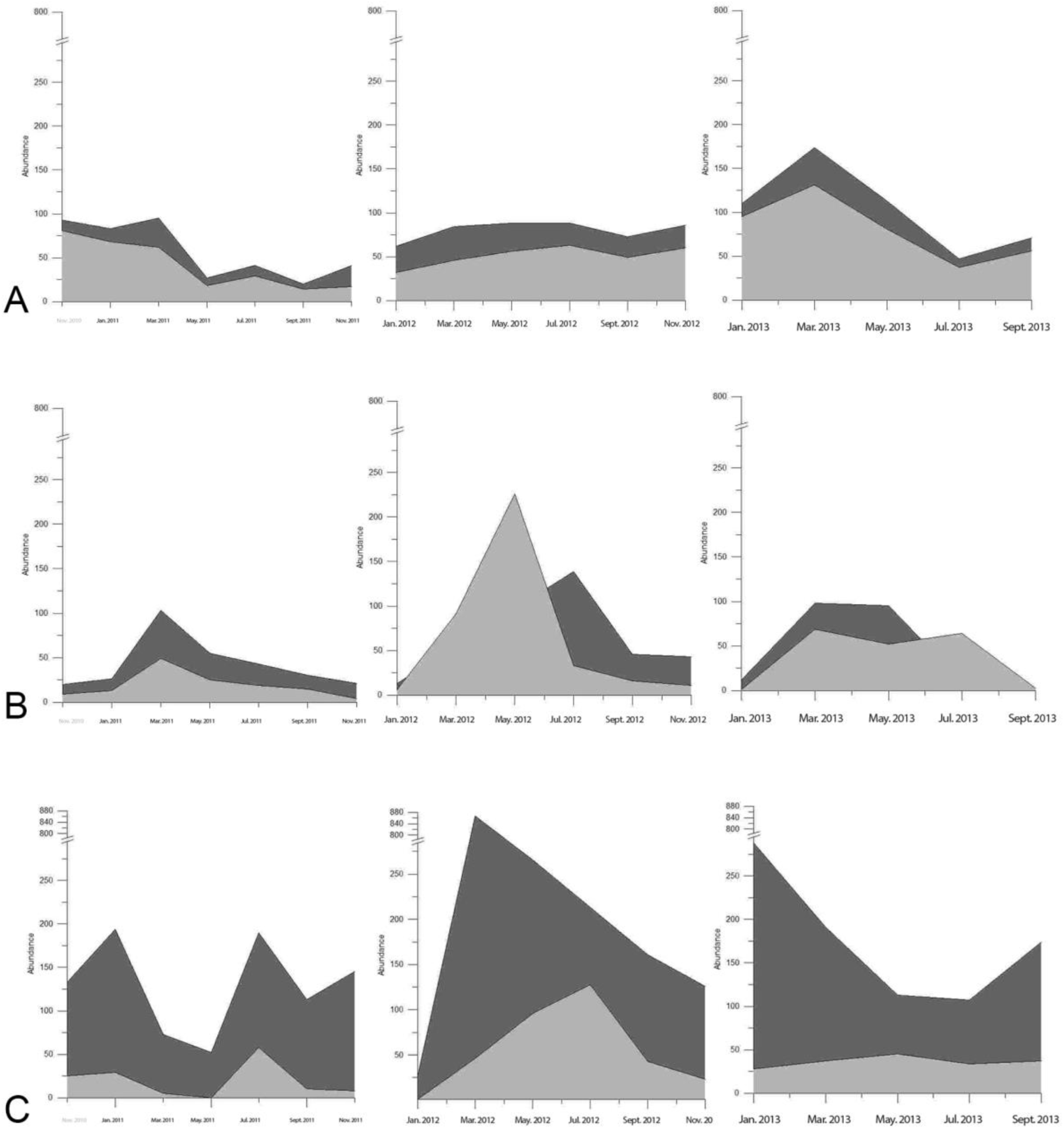
Monthly butterfly abundance recorded across study sites and years in Western Ecuador from Nov 2010 to Sept 2013. A) Wet forest, B) Transition forest and C) Dry forest.

In general, both intra- and inter-annual variation in butterfly abundance was correlated with rainfall in seasonal forests but not in the aseasonal forest. For example, butterfly communities from the southern seasonal forests in Manabi and Loja showed highest numbers of species and individuals in 2012, coinciding with the wettest year covered by the sampling period (cumulative rain per year equaled 1763 mm^3^ and 1203 mm^3^, respectively). Similarly, rain was considerably higher during 2012 in the wet forest (3886.43 mm^3^), nevertheless the most conspicuous butterfly peak occurred one year later (*i.e.*, nearly three times more species and individuals sampled compared to previous years). In addition, this pattern remained even when splitting the butterfly community into wet forest and dry forest species (Fig. 4). Peak abundance of dry forest species was synchronized in both seasonal forests of Manabi and Loja through 2012, matching the year with highest water availability in these ecosystems (Fig. 1). On the contrary, the abundance of wet-forest species reached a peak during the wettest year in the Manabi transition forest, whereas this peak showed a 1-year time lag after the wettest year in the Canande wet forest (Fig. 4).

**Figure 4.**
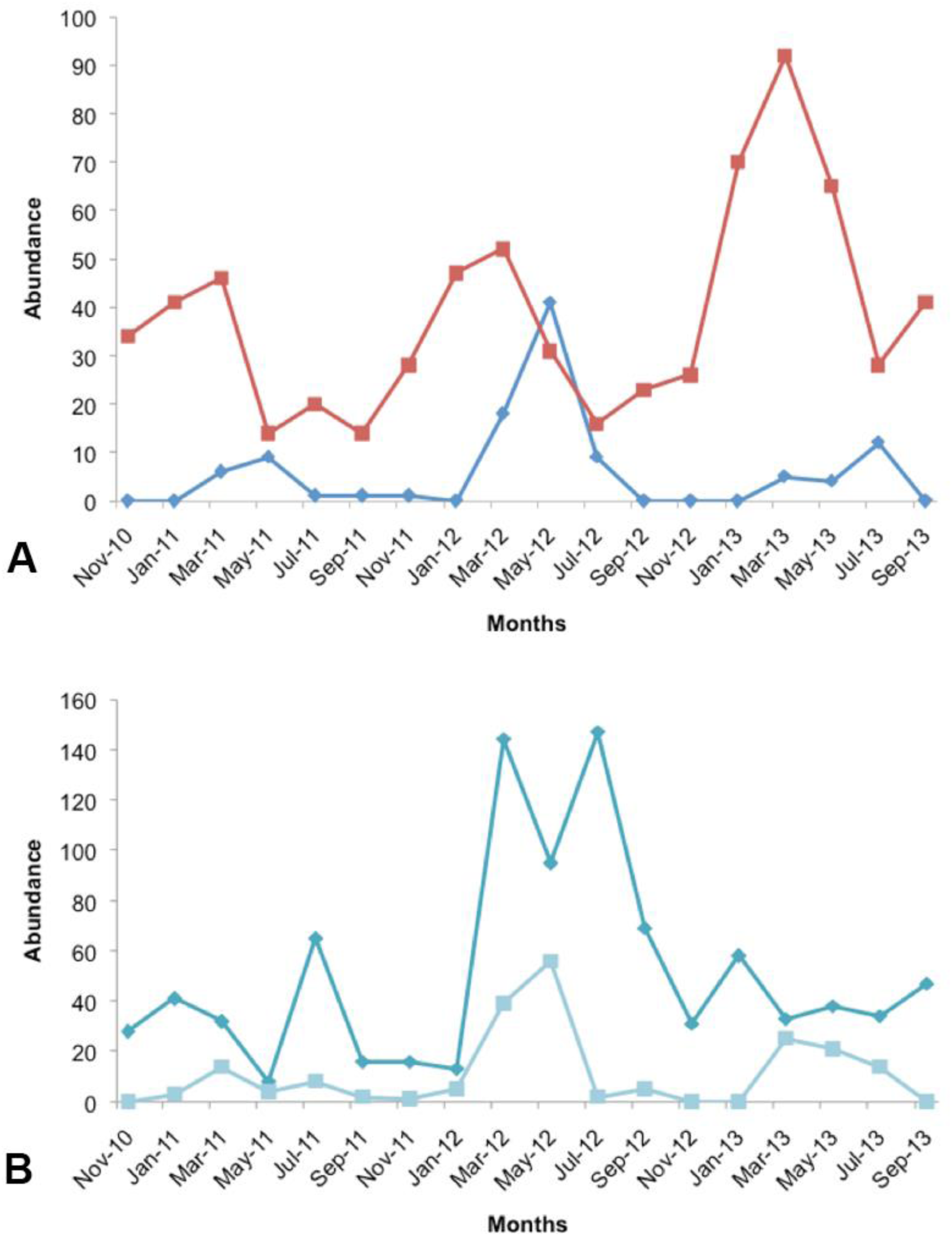
Monthly abundance of wet and dry species recorded across study sites from Nov 2010 to Sep 2013. A) Wet species recorded in the wet forest (red line with squares) and transition forest (blue line with triangles), B) Dry species registered in the transition forest (line with squares) and dry forest (line with triangles).

For some of the most common species, occurring in all three sites, the highest number of individuals occurred at the same time of year across all sites and years (Fig. 5). For example, the highest peak in abundance of *H. amphinome* coincided across ecosystems through the 2012 wet season but extended towards the driest part of the year in the wet forest. Conversely, *F. ryphea* exhibited the same highest peak across years only in the seasonal forests, peaking through the 2012 wet season, but showed a 1-year time lag in the wet forest. For other species, there was not an obvious pattern; the population abundance of *S. blomfildia* peaked distinctively across study sites (Fig. 5).

**Figure 5.**
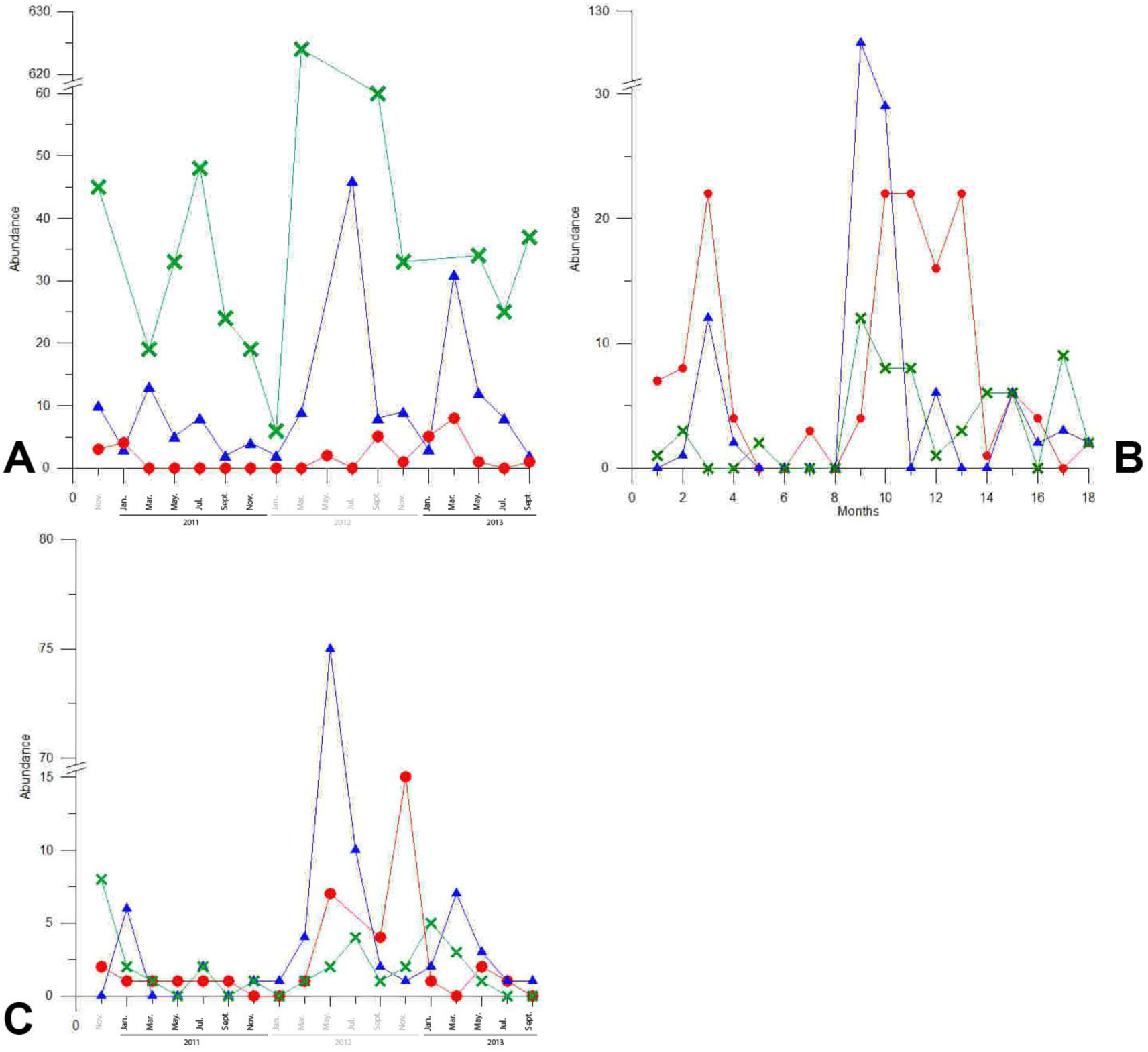
Monthly abundance of species distributed in all three study sites: wet forest (red line), transition forest (blue line) and dry forest (green line). A) *Fountainea ryphea*, B) *Hamadryas amphinome* and C) *Smyrna blomfildia.*

Generalized linear models revealed that rainfall was significantly related to monthly butterfly abundance over all ecosystems, and to monthly species richness only in the wet forest (Table 1). Furthermore, rainfall coefficients were positive, indicating increased cumulative monthly rainfall is associated with higher butterfly numbers. Temperate was not a significant predictor in all models, except when species richness was analyzed for the wet and transition forest (estimate= −0.08, p <0.05, estimate= −0.34, p <0.05, respectively), and in contrast to rainfall, the temperature coefficient was negative indicating that warmest months had lower species richness.

**Table 1.** Results of GLM per each study site (*i.e.*, wet, transition and dry forests) using monthly abundance or observed species richness as the response variable and monthly measures of climatic variables as predictors. Models included the negative binomial distribution for the response variables. Asterisks and double asterisks represent significant and highly significant associations between butterfly numbers and climate, respectively.

| Predictors | Abundance<br>Coefficients | *t* value | P | Species<br>Coefficients | richness<br>*t* value | P |
| --- | --- | --- | --- | --- | --- | --- |
| Wet forest |  |  |  |  |  |  |
| Total rain | 0.001 | 2.48 | 0.01* | 0.00 | 2.97 | 0.00** |
| Mean temperature | -0.10 | -1.72 | 0.09 | -0.08 | -2.15 | 0.03* |
| Transition forest |  |  |  |  |  |  |
| Total rain | 0.01 | 3.08 | 0.00** | 0.00 | 1.29 | 0.20 |
| Mean temperature | -0.34 | -1.39 | 0.17 | -0.34 | -2.16 | 0.03 * |
| Dry forest |  |  |  |  |  |  |
| Total rain | 0.00 | 3.01 | 0.00** | 0.00 | 0.74 | 0.46 |
| Mean temperature | -0.09 | -0.62 | 0.54 | -0.16 | -1.50 | 0.13 |

Similarly, total monthly rainfall, but not temperature, was significantly negatively associated with butterfly species seasonality in the dry forest (Estimate= −0.13, p=0.00) (Table 2). These results suggest butterfly species with stronger seasonality as measured by the CV of their relative monthly abundance were significantly more constrained by rainfall variation. Among more seasonal species is *Ithomia cleora* (CV= 3.65), a range-restricted species confined to western Ecuador and northwest Peru, mostly sampled in May 2012 (6 out of 7 individuals).

**Table 2.** Results of GLM for the dry forest communities using the coefficient of variation of butterfly abundance (measure of seasonality) as dependent variable and climatic variables as predictors. The response variable better fit the normal distribution. Double asterisks represent rainfall highly significantly affects butterfly seasonality.

| Predictors | Coefficients | t value | P |
| --- | --- | --- | --- |
| Rain (mm3) | -0.13 | -3.63 | 0.00** |
| Std.Temperature (°C) | 0.39 | 0.68 | 0.50 |
| Std.Relative Humidity (%) | -0.21 | -1.04 | 0.30 |

## Discussion

Butterfly communities exhibited conspicuous intra and inter-annual variation in temporal dynamics with certain elements (*e.g.*, maximum abundance recorded) of seasonality patterns likely synchronized in seasonal forests across years but not in aseasonal forests. In addition, the highest numbers of species and individuals occurred during the wet season across all study sites and years; indeed, rainfall was significantly positively associated with temporal abundance. Likewise, butterfly species displaying stronger seasonality were significantly associated with higher rainfall periods in seasonal forests. As far as I know, this represents the first study that simultaneously sampled and described seasonality patterns of tropical butterfly communities across ecosystems with varying climatic seasonality, including data emphasizing variation in precipitation regimes might significantly affect more seasonal species.

The pattern of butterfly abundance peaking through the wet season found here is concordant with the majority of studies into temporal dynamics of tropical insects (Wolda 1978, Novotny & Basset 1998, Grimbacher *et al.* 2009, for a review Kishimoto-Yamada & Itioka 2015), and particularly with studies focused on butterfly communities distributed in Neotropical dry forests (Shahabuddin *et al.* 1999, Pozo *et al.* 2008, Torres *et al.* 2009, Checa *et al.* 2010, Checa 2014). Precipitation, but not temperature, was significantly associated with butterfly abundance across ecosystems. By contrast, temperature and species richness were significantly related in the wet forest, a result concordant with previous research in rainforest butterflies (Checa *et al.* 2009, Ribeiro *et* al. 2010, Grøtan et al. 2014). This result could be explained by the presence of different limiting climatic factors in seasonal dry forests compared to aseasonal wet forests, and the associated phenology of the flora and fauna. Maintaining a sufficiently high thoracic temperature for adequate flight performance is a greater problem for butterflies inhabiting wet forests compared to butterfly species from drier habitats, where water availability instead is more critical.

The close relationship between butterfly numbers and climatic variables was expected because butterflies are ectothermic and their thermal physiology is thus highly dependent on climate (Roy *et al.* 2001). Weather affects population growth by limiting the available time for adult flight needed for oviposition (Warren 1992) and by its effects on survival of the immature stages (Warren 1992, Hellmann 2002, Dooley *et al.* 2013) and fecundity (Boggs & Freeman 2005). Other factors are also important such as predator avoidance, courtship and adult feeding (see below).

Higher availability of food sources for adult butterflies (*i.e.*, fruits, carrion) and larvae (leaves) could explain the peaks in butterfly numbers during the wet season, in addition to the significant association of rainfall with butterfly numbers since water availability is more critical in drier habitats and seasons where dehydration of both hostplants and butterflies is at risk. Fruiting peaks and increased leaf production occur during the rainy season in tropical forests (Foster 1996 in Grøtan *et al.* 2012). Indeed, weather-related density-independent processes (*e.g.*, senescence timing of hostplants) are considered the main driver of population dynamics for some temperate butterfly species (Hellman *et al.* 2004, Yamamoto *et al.* 2007), and insect phenology thus follows and synchronizes with host-plant phenology, which also varies from year to year in response to environmental conditions (van Asch & Visser 2007, Valtonen *et al.* 2013). Additionally, the wet season is likely the main period of oviposition (Shahabuddin *et al.* 1999) since leaf-feeding butterflies prefer new shoots and leaves for this task (Rodrigues & Pires 1999, van Asch & Visser 2007, Srygley *et al.* 2010). In the seasonal forests, the abundance peak of adults in March, a few months after the start of the rainy season in December, could result from the time-lag due to the development of the immature stages. As climate change has affected the phenology of temperate butterflies (Roy & Sparks 2000, Stefanescu *et al.* 2003, Altermatt 2010), disrupting the relationship between butterflies and their hostplants and adult resources, similar consequences can be expected in dry forests owing to the close relationship between rainfall and the temporal dynamics of their butterfly communities.

The abundance of adult butterflies does not necessarily reflect total population abundance, since individuals may be present at a site but undetected as diapausing adults or immature stages (Kishimoto-Yamada & Itioka 2015). Diapause, a state of low metabolic activity prevents development during unfavorable conditions increasing resistance to adverse environmental conditions (van Asch & Visser 2007). It is possible butterflies survive the dry season as adults in reproductive diapause (see Shahabuddin *et al.* 1999), although Lepidoptera can also diapause as pupae, eggs or more rarely larvae (Torres *et al.* 2009).

Species detection probability or detectability may additionally bias the observed seasonal patterns. For example, sunny days enhance butterfly activity, hence increasing probability of trap captures, and the level of bait attractiveness may differ over seasons (Torres *et al.* 2009, Ribeiro et al. 2015). Although these factors can certainly explain some variation, the bias produced is likely to be negligible in this study since butterfly abundance peaks were detected during the wettest part of the year (the dry season encompasses a larger number of sunny days which increases the probability of both trap captures and scheduling sampling during warmer days), and differences in abundance among years and seasons were relatively large (see Checa *et al.* 2014). Additionally, in Neotropical forests, detectability of butterfly species attracted to baits has been reported to positively correlate with observed abundance, which means probability detection raises as butterfly abundance increases (Ribeiro *et al.* 2015).

Peaks in abundance and species richness has been reported for rain forests during the transitional months rather than throughout the wet season in Costa Rica (Grøtan *et al.* 2014) and Amazonia (see DeVries & Walla 2001, Checa *et al.* 2009, Grøtan *et al.* 2012), results that contrast with those found here for wet forests, where butterfly numbers reached a peak during the most humid part of the year. These differences are likely related to species detectability and underlying factors regulating temporal dynamics and assembly mechanisms shaping communities, and how they vary across space (*i.e.*, biogeography regions, biomes) (see below).

Similarities were detected in butterfly community dynamics across the two seasonal sites, in particular with both showing the greatest butterfly abundance during the 2012 wet season, which coincided with highest rainfall recorded across years and ecosystems. Meanwhile in the aseasonal wet forest, the highest annual abundance occurred one year later. This pattern may indicate that regional processes are more prevalent in regulating butterfly populations in dry forests (*e.g.*, climatic variables) compared to local mechanisms (*e.g.*, predation, natural enemies), which may be more prevalent drivers of temporal patterns in the wet forest. Further evidence to support this idea of variable drivers of community dynamics comes from varying seasonal patterns of wet/dry species depending on the sampled ecosystem. The highest abundance for dry forest species was synchronized in both seasonal forests in 2012, whereas wet forest species also reached a peak during the wettest year in the Manabi transition forest, but one year later in the wet forest. In addition, different regulating mechanisms may operate at the species level. Regional climatic factors might be more relevant for *H. amphinome* populations since its abundance peak coincided across ecosystems through the 2012 wet season. On the other hand, although *F. ryphea* also reached its peak annual abundance in 2012 during the wet season within the seasonal forests, it reached peak abundance one year later in the wet forest, which may perhaps further indicate the importance of local processes for population regulation.

In temperate areas, the synchrony of population dynamics among species can be explained by a strong effect of environmental forces (Nowicki *et al.* 2009). In England, butterfly populations are also often regionally synchronized (see Pollard 1991) due to the regional correlation in climatic patterns (Sutcliffe *et al.* 1996). Synchrony occurs because species show similar responses to environment (*i.e.*, positive covariation) with intra- or inter-specific interactions playing a weak role in shaping temporal abundance patterns of species within a community (Mutshinda *et al.* 2010).

Furthermore, time-lags of peaks in abundance in response to weather conditions occur particularly if different stages of the life cycle are affected in overlapping populations (Boggs & Inouye 2012). There is a frequent correlation between increased humidity/precipitation in one year and increased butterfly abundance in the following year (Pollard 1991, Forister *et al.* 2011, Dooley *et al.* 2013). In one study, the mechanisms involved included an increase in nectar availability through higher precipitation, which in turn enhanced fecundity (O’Brien *et al.* 2004); but precipitation can also depress natural enemies and permit increased growth rates in the population (Forister *et al.* 2011). Therefore, weather conditions can lead to density-dependent effects the following year. Positive effects of temperature during one previous year on growth rates of the following year have also been reported (Roy *et al.* 2001, Forister *et al.* 2011, WallisDeVries *et al.* 2011). These effects may occur because higher temperatures can extend flight periods and hence permit more oviposition opportunities, or warmer environments may enhance plant growth supporting higher larval survival.

Predators and parasitism are considered major regulatory factors of tropical butterflies (Gilbert 1972), with, for example, selection from bird predators (*e.g.*, jacamars, flycatchers, kingbirds) believed to be responsible for the remarkable mimicry observed in butterflies (Langham 2004). Nevertheless, our understanding of the importance of competition, predation and parasitoids in limiting butterfly abundance is very limited (Grøtan *et al.* 2012). In studies focused on population dynamics, the detection of density-dependence is greatly limited by short-term abundance studies (Woiwood & Hanksi 1992). Therefore, a lack of detection of density dependence can result from short times series analyzed (<20 years) rather than weak contributions to population regulation (Mittelbach 2012). Moreover, some authors argue that density-dependence has been underestimated in butterflies due to a greater attention on the adult stage (Nowicky *et al.* 2009). As a consequence of the long-term abundance data required and the need for monitoring factors shaping fluctuations, very few studies have quantitatively analyzed the relative impact of density-dependent factors and density-independent factors in the population regulation of butterflies, and the few studies that have were focused on very well-known species from temperate areas (Azerefegne *et al.* 2001, Mclaughlin *et al.* 2002, Nowicky *et al.* 2009, Dooley *et al.* 2013). Further research is therefore needed on density-dependent factors (*e.g.*, competition, predation, parasitism) to have a more complete understanding of the population dynamics of tropical butterflies and to verify what has been suggested here, that local density-dependent mechanisms (e.g, predation, competition) are more significant in controlling abundance in wet forests compared to dry forests.

Our understanding of mechanisms influencing seasonal patterns and the evolution of insect seasonality might also benefit from research at the species level, taking into account generation cycles, different life stages and specific behaviors such as diapause (see Kishimoto-Yamada & Itioka 2015). In addition studies are needed of the variation of populations across microhabitats and geographic space, since the relative importance of density-dependence or density independence for butterfly populations can vary across space and time (Flockhart *et al.* 2012, Dooley *et al.* 2013). In terms of distribution, temperate butterfly populations show greater fluctuations at the edges of their distributions (Warren 1992; Thomas *et al.* 1994), and are more strongly affected by environmental stochasticity compared to populations occurring in the center (Nowicky *et al.* 2009). At very local spatial scales, in adjacent habitats, different factors become increasingly important; butterflies from homogenous habitats may be more influenced by weather (*i.e.*, rainfall in the preceding year) rather than by endogenous factors (McLaughlin *et al.* 2002).

Finally, a significant result found in the present study is that butterflies with stronger seasonality were significantly negatively associated with total monthly rainfall but not variability of both temperature and relative humidity in seasonal forests. As a consequence, butterfly species with uneven temporal distribution (*i.e.*, restricted ecological niche) might be more influenced by changes in the precipitation regime within forests with marked climatic seasons. It is likely some strongly seasonal species may also be restricted in geographic range, as is the case with *Ithomia cleora*, which could amplify its susceptibility to climate change if effects of rainfall variation prove to be negative (*i.e.*, narrowing even more peaks of occurrence). Hence, seasonality might be a further trait contributing to extinction vulnerability of taxa, along with other previously studied traits, including body size (Brown 1971, Kattan *et al.* 1994, Shahabuddin & Ponte 2005), population density (Pimm *et al.* 1988, Bolger *et al.* 1991, Shahabuddin & Ponte 2005) and mobility (Graves & Gotelli 1983). In addition, more seasonal nymphalid species tend to also have larger body sizes (Ribeiro & Freitas 2011), a trait related to increased vulnerability to extinction in some Neotropical nymphalids (Shahabuddin & Ponte 2005). Further research about this topic will shed more light into whether seasonality is another contributing factor for extinction vulnerability of insect species.

The results of this study are also relevant for determining the possible impacts of climate change on tropical ectotherms, which several authors regard as potentially more sensitive than temperate organisms (Wright *et al.* 2009, Laurance *et al.* 2011) due to their narrower thermal tolerance compared to temperate species (Deutsch *et al.* 2008). Further research into insect seasonality taking into account species traits such as host specialization or geographic distribution, are needed to achieve that goal. For example, such studies might shed light on how insects may adjust to unfavorable seasons when extending their distribution to higher altitudes (Kishimoto-Yamada & Itioka 2015), or provide information about the climatic niche of species at local scales, which in turn can further contribute to understanding species distribution and latitudinal patterns of species richness. This type of research is a high priority for the fragmented dry forests in Ecuador as climate change is a major threat because of the positive feedback between forest fragmentation and drought (Laurance & Williamson 2001). Moreover, these fragmented forests are more vulnerable to the impacts of El Nino-Southern Oscillation (ENSO) droughts, whose frequency and therefore impact on butterfly ecology (Srygley *et al.* 2010) is expected to be altered by climate change, in addition to an increased variability of weather patterns (Easterling *et al.* 2000).

